# Multiple fertility restorer loci for cytoplasmic male sterility caused by *orf137* in tomato

**DOI:** 10.1101/2025.01.31.635880

**Authors:** Yurie Iki, Issei Harada, Kentaro Ezura, Seira Mashita, Kosuke Kuwabara, Hitomi Takei, Atsushi Toyoda, Kenta Shirasawa, Tohru Ariizumi

**Affiliations:** Graduate School of Life and Environmental Sciences, University of Tsukuba, Tsukuba, Ibaraki 305-8572, Japan; Institute of Life and Environmental Sciences, University of Tsukuba, Tsukuba, Ibaraki 305-8572, Japan; Advanced Genomics Center, National Institute of Genetics, Mishima, Shizuoka 411-0801, Japan; Department of Frontier Research and Development, Kazusa DNA Research Institute, Kisarazu, Chiba 292-0818, Japan; Tsukuba Plant Innovation Research Center, University of Tsukuba, Tsukuba, Ibaraki 305-8572, Japan

**Keywords:** Cytoplasmic male sterility, F_1_ seed production, Fertility restoration, Genome, Tomato, Transformation

## Abstract

Cytoplasmic male sterility (CMS) in plants is caused by incompatibility between nuclear and cytoplasmic genetic information. Fertility can be restored through the action of fertility restoration (*RF*) genes, which are usually present in the nucleus. CMS lines of tomato (*Solanum lycopersicum*) have been developed from asymmetric cell fusions, in these lines, cultivated tomato served as a nuclear donor and its wild relative, *S. acaule*, as a cytoplasm donor. Although *RF* genes are present in wild relatives of tomato, no genetic or genomic information on the RF genes is yet available. This study reports an *RF* genetic locus, *RF1*, on chromosome 1 of *S. pimpinellifolium* LA1670 and *S. lycopersicum* var. *cerasiforme* LA1673 that was revealed by bulked segregant analysis and sequencing. An additional *RF* locus, *RF2*, was identified on chromosome 2 of LA1670. A transgenic approach identified two other candidate genes that also restored fertility. The genomic sequence of *S. cheesmaniae* LA0166 was assembled using high-fidelity, long-read sequencing technology. Sequence comparisons identified further candidate *RF* genes on chromosome 1 of *S. cheesmaniae* LA0166. These results suggested that multiple gene loci control the fertility restoration trait in wild relatives of tomato.

## Introduction

Cultivated and wild species have evolved separately and so, even when two species are cross-compatible, their genetic information may be incompatible between species (Orr, 1996). Following the generation of an interspecific hybrid, recurrent backcrossing, intended to replace a wild nuclear gene with a cultivated one, or *vice versa*, can lead to male sterility due to incompatible interactions between the nuclear and cytoplasmic genomes. This is known as cytoplasmic male sterility (CMS). CMS occurs when plants lack nuclear genes able to regulate mitochondrial gene expression, thus exposing the deleterious effects of mitochondrial genes (Toriyama *et al*., 2024). As fertility can be restored by nuclear genes that are compatible with the mitochondrial genes, the gene regulators encoded in the nuclear genome are called fertility restoration (*RF*) genes.

CMS lines of tomato (*Solanum lycopersicum*), which is one of the most important food crops worldwide, were developed by asymmetric cell fusions that used cultivated tomato as a nuclear donor and its wild relative, *S. acaule*, as a cytoplasm donor (Melchers *et al*., 1992). A mitochondrial gene, *orf137*, triggers CMS in the fusion lines (Kuwabara *et al*., 2022). *S. pimpinellifolium* LA1670, *S. lycopersicum* var. *cerasiforme* LA1673, and *S. cheesmaniae* LA0166, three wild relatives of tomato, possess *RF* genes that restore fertility to the CMS lines (Harada *et al*., 2003); to date, however, no *RF* genetic loci have been reported in tomato.

CMS is a useful trait for F_1_ seed production because emasculation, which is labor-intensive and thus costly, is not required for outcrossing. In crops whose product is set fruit and seeds, however, the paternal lines of F_1_ hybrids must possess *RF* genes to ensure fertility in their F_1_ progeny, whose cytoplasm contains CMS genes. Identification of *RF* gene loci is thus an important aspect of efficient breeding in the paternal lines. Pentatricopeptide repeat (PPR) proteins act as *RF* genes in many crop plants (Toriyama, 2021). These genes are often tandemly arrayed in *RF* regions of genomes (Geddy and Brown, 2007); it is also known that the number of copies of *PPR* genes varies between *RF* loci and that this copy number variation functions in fertility restoration (Wang *et al*., 2006).

As the genomes of non-RF tomato lines may lack functional *RF* gene sequences, the fully sequenced genomes from RF lines are useful tools in the identification of *RF* genes. The genomic sequences of *S. pimpinellifolium* LA1670 and *S. lycopersicum* var. *cerasiforme* LA1673, two of the three relatives of tomato that possess *RF* genes, are publicly available (Takei *et al*., 2021), although the genome of the third wild relative, *S. cheesmaniae* LA0166, is not. In this study, we used the genomic sequences of the RF lines to identify genetic loci for *RF* genes in all three tomato relatives. Furthermore, we determined the genomic sequence of *S. cheesmaniae* LA0166. Finding the locations of these *RF* genes, together with the development of linked DNA markers, is a major step towards the identification of *RF* genes in tomato that will enable the efficient breeding of elite RF paternal lines for use in tomato F_1_ seed production.

## Materials and Methods

### Plant materials

The full details of the pedigrees of the tomato lines used in this study are shown in Supplementary Figure S1. The tomato (*Solanum lysopersicum*) cultivars O and Micro-Tom, both of which lack CMS genes in the cytoplasm and *RF* genes in the nucleus, were used in experiments. We also used the CMS tomato lines CMS[MSA1] and CMS[P] that were described in our previous study (Kuwabara *et al*., 2021), CMS[MSA1] was developed by repeated backcrossing using O as a paternal recurrent parent and the maternal male-sterile line MSA1 as a cytoplasmic donor; MSA1 is an asymmetric cell fusion line derived from the cultivar Sekai-ichi (nuclear donor) and a wild relative *S. acaule* (cytoplasmic donor) (Melchers *et al*., 1992). CMS[P] was produced by backcrosses in which the cultivar P (Kuwabara *et al*., 2021) served as the paternal recurrent parent and an asymmetric cell fusion between P (nuclear donor) and *S. acaule* (cytoplasmic donor) as the maternal line (Kuwabara *et al*., 2021). Both CMS lines carry the CMS gene, *orf137*, in their mitochondria (Kuwabara *et al*., 2022). In addition, we used Dwarf CMS[P], a line developed by repeatedly backcrossing CMS[P] with Micro-Tom (TOMJPF0001) (Kuwabara *et al*., 2021).

The three wild relatives of tomato, *S. pimpinellifolium* LA1670, *S. lycopersicum* var. *cerasiforme* LA1673, and *S. cheesmaniae* LA0166, were used as RF lines. Seeds from these species were obtained from the Tomato Genetic Resource Center at the University of California, Davis, USA. All plants were grown in greenhouses at the University of Tsukuba for phenotype evaluation.

### Pollen germination test

Phenotyping of pollen tube growth was performed using the method described in Kuwabara (2022). For *in vivo* pollen germination tests on stigmas, pistils were fixed in ethanol: acetic acid (3: 1, v/v) 24 h after self-pollination. Fixed pistils were soaked in 5 M NaOH for 24 h. After washing three times, pistils were stained in the dark for 24 h with 0.001 w/v% aniline blue in 0.1 M K_2_HPO_4_ buffer (pH 10).

Pollen germination tests using liquid germination medium were performed according to a previously described protocol with minor modifications (Steven and Wouter, 1993). Freshly opened flowers were soaked in 1 ml germination medium [15.1 w/v% polyethylene glycol (average molecular weight: 6000), 10 w/v% sucrose, 1.63 mM H_3_BO_3_, 1.27 mM Ca(NO_3_)_2_, 1 mM MgSO_4_, 1 mM KNO_3_, and 0.1 mM K_2_HPO_4_]. The mixture was strongly vortexed to release the pollen from the anthers. The flower residues were removed and the pollen suspension was incubated in the germination medium and agitated using a rotator at 25°C for 4 h. Pollen tubes were observed using a BX53 microscope (Olympus, Hachioji, Japan). Digital images were captured using a DP72 camera (Olympus).

### DNA extraction and DNA marker analysis

Genomic DNA was isolated from young leaves using a DNeasy Plant Mini Kit (Qiagen, Hilden, Germany) and used as a template for PCR. Each 10 µl PCR contained 0.5□µl genomic DNA, 0.3□µM primers (Supplementary Table S1), 2× PCR buffer (TOYOBO, Osaka, Japan), 400□µM dNTPs, and 1 U DNA polymerase (KOD FX Neo, TOYOBO). The thermal cycling conditions were as follows: initial denaturation at 94°C for 3□min; 35 cycles of denaturation at 98°C for 15□s, annealing at 55-60°C for 30□s, and extension at 68°C for 60□s; followed by a final extension at 68°C for 3□min. The PCR products were digested with restriction enzymes and separated by electrophoresis through 1% agarose gels submerged in Tris-acetate-EDTA (TAE) buffer. Gels were stained with Midori Green Advance (NIPPON Genetics, Tokyo, Japan) to visualize DNA bands under ultraviolet illumination.

### Bulked segregant analysis with short-read sequencing

A whole genome shotgun library was prepared using TruSeq DNA PCR-Free Sample Prep Kit (Illumina, San Diego, CA, USA). A ddRAD-seq library was constructed as described previously (Shirasawa *et al*., 2016) with minor modifications. Genomic DNA was digested with the restriction enzymes, *Pst*I and *Msp*I (Fast Digest restriction enzymes; Thermo Fisher Scientific, Waltham, MA, USA). Nucleotide sequences were determined using short-read DNA sequencers manufactured by Illumina and MGI Tech (Shenzhen, China). Data processing and single nucleotide polymorphism (SNP) identification were performed as described previously (Shirasawa *et al*., 2016). High-quality sequence reads were mapped onto the reference sequence to identify high-confidence SNPs. SNPs associated with the traits were identified and visualized using QTLseqr (Mansfeld and Grumet, 2018).

### RNA-Seq analysis

Total RNA was extracted from anthers and pollen grains using an RNeasy Plant Mini Kit (Qiagen) and treated with RNase-free DNase (Qiagen). Sequence libraries were prepared from total RNA using a TruSeq Stranded mRNA Library Prep Kit (Illumina) and random hexamers as primers for reverse transcription. The resulting libraries were sequenced on a DNBSEQ G400 instrument (MGI Tech) to generate 100-bp paired-end reads. The obtained reads were mapped onto the LA1670 genome using HISAT2; StringTie was used to quantify expression by tags per million score (Pertea *et al*., 2016). Mitochondrial target proteins were predicted using MitoFates (Fukasawa *et al*., 2015), Deepmito (Savojardo *et al*., 2020), and TargetP (Emanuelsson *et al*., 2000).

### Genetic transformation

To construct an expression vector for *RF* genes, the DNA sequences encoding the genes *SPI01g07345* and *SPI01g0748*5 plus 1 kb of upstream sequence as a promoter were artificially synthesized using gene synthesis technology (Azenta Life Sciences, Chelmsford, MI, USA) and ligated into the binary vector pBI101. The plasmids were introduced into the Dwarf CMS[P] line using *Agrobacterium*-mediated transformation, as described by (Sun *et al*., 2006). Transformants were selected using kanamycin resistance and subjected to pollen germination tests.

### Genome assembly and gene annotation of LA0166

LA0166 genome assembly was performed as described previously (Shirasawa and Ariizumi, 2024). High molecular weight genomic DNA was isolated from leaves of LA0166 using Genomic-Tips (Qiagen). gDNA libraries were constructed using the SMRTbell Express Template Prep Kit 2.0 (PacBio, Menlo Park, CA, USA). The resulting libraries were fractionated with BluePippin (Sage Science, Beverly, MA, USA) to eliminate fragments shorter than 20 kb and sequenced on a SMRT Cell 8M using the Sequel II system (PacBio). HiFi reads were constructed using the CCS pipeline (https://ccs.how) and assembled using Hifiasm with default parameters. The assembly was aligned on the SL4.0 genomic sequence (Tomato Genome Consortium, 2012) to establish a chromosome-level sequence with RaGoo (Alonge *et al*., 2019).

Gene prediction was performed using BRAKER3 (Gabriel *et al*., 2024), based on the peptide sequences of the genes predicted by ITAG4.0 (https://solgenomics.net) and Iso-Seq reads obtained from anthers of LA0166, according to the PacBio standard protocol (PacBio). The gene sequences reported by ITAG4.0 were then mapped onto the pseudomolecule sequences using Liftoff (Shumate and Salzberg, 2020). The completeness of the genome assembly and gene prediction were evaluated with embryophyta_odb10 data using benchmarking universal single-copy orthologs (BUSCO) (Simão *et al*., 2015). Repetitive sequences in the assembly were predicted by RepeatMasker (https://www.repeatmasker.org) using repeat sequences registered in Repbase and a *de novo* repeat library was built using RepeatModeler (https://www.repeatmasker.org). Genome structures were compared with MCScanX (Wang *et al*., 2012) and visualized using SynVisio (Bandi and Gutwin, 2020).

## Results

### *RF1a1 in* S. pimpinellifolium *LA1670*

The mode of fertility restoration in *S. pimpinellifolium*, LA1670, was investigated using an F_2_ population (n = 368) derived from the cross CMS[MSA1] × LA1670 (Supplementary Figure S1A). All but five F_2_ plants achieved fruit set by self-pollination. Pollen from the five plants that did not set fruit, or set seedless fruits due to parthenocarpy, was able to germinate, however, suggesting that the gametophytic mode of pollen fertility restoration was present in LA1670.

Subsequently, a BC1F1 population (n = 244) was generated by crossing an F_1_ hybrid (derived from CMS[MSA1] × LA1670) with O (Supplementary Figure S1A). When the BC1F1 population had grown to maturity, 133 plants showed seed formation and/or pollen germination, whereas the remaining 113 plants exhibited no pollen germination, indicating male sterility. The segregation ratio of fertile to sterile plants was close to 1:1 (χ^2^ = 1.328, *P* = 0.249), suggesting that fertility restoration was controlled by a single genetic locus.

To identify the genetic locus underlying fertility restoration, we performed bulked segregant analysis by sequencing. Sequence reads, obtained from bulked samples of the fertile and sterile BC1F1 plants, were mapped onto the genomic sequence of *S. pimpinellifolium*, LA1670. This analysis suggested that an RF candidate gene was located in a 10-Mb region situated between approximately 80 and 90 Mb of chromosome 1 (Supplementary Figure S2). Next, 46 BC3F1 plants, generated by backcrosses between a fertile BC1F1 plant as a donor and O as a recurrent parent (Supplementary Figure S1A), were genotyped using 31 cleaved amplified polymorphic sequence (CAPS) markers designed against the candidate region. This analysis narrowed the candidate region down to a 2.4-Mb region between 82.6 and 85.0 Mb of chromosome 1 (Figure 1). This region was named *RF1a1*.

**Figure 1.**
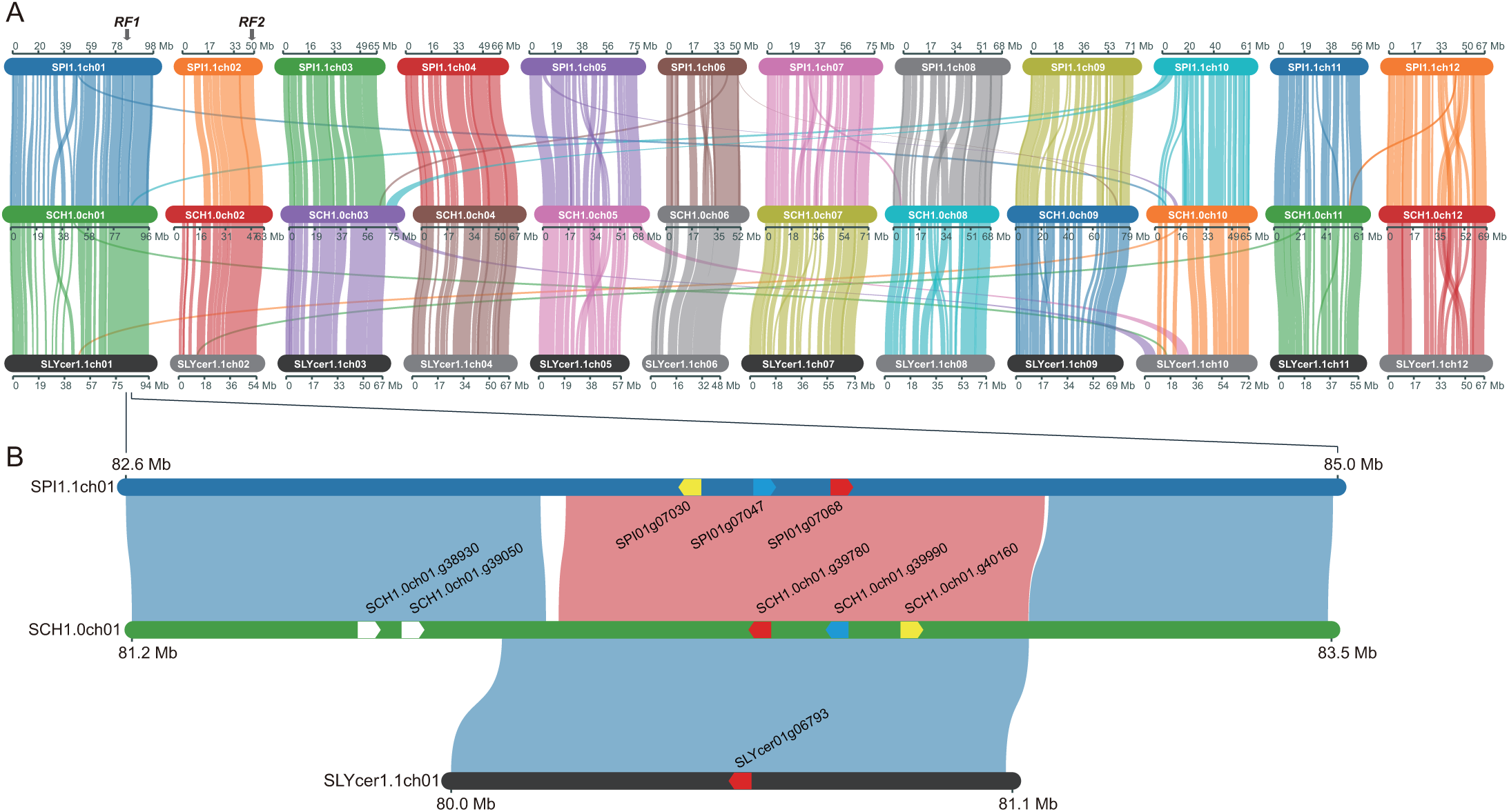
Genomic structures of the LA1670, LA0166, and LA1673 RF lines. A. Chromosome structural similarity between the three RF lines of LA1670 (SPI1.1ch01 to SPI1.1ch12), LA0166 (SCH1.0ch01 to SCH1.0ch12), and LA1673 (SLYcer1.1ch01 to SLYcer1.1ch12). Arrows indicate the RF1 and RF2 loci. B. Genomic structural similarity between the RF1 loci on chromosome 1 of the three RF lines. Blue and red alignment links between the RF lines show regular and reversed orientations, respectively. Pentagons indicate *PPR* genes with gene ID colors (yellow, blue, and red) indicating orthologous genes.

### RF1b and RF1c in S. pimpinellifolium LA1670

As some tandemly repeated *PPR* genes at *RF* loci restore male fertility, we also surveyed *PPR* genes outside the candidate region. This analysis identified 13 *PPR* genes (Supplementary Table S2), of which two, *SPI01g07345* and *SPI01g07485*, satisfied our screening criteria, as they 1) were transcribed in anthers and/or pollens; 2) possessed missense mutations between fertile and sterile lines; and 3) were predicted to be mitochondrial target proteins. We introduced these genes separately into Dwarf CMS[P] to produce transgenic lines. *SPI01g07345* restored male fertility in one of the 22 independent transgenic lines (Figure 2A). Likewise, *SPI01g07485* also restored male fertility in one of the 22 independent transgenic lines (Figure 2B). These results indicated that both genes could restore fertility in CMS tomato lines. We therefore renamed *SPI01g0734* and *SPI01g0748* as *RF1b* and *RF1c*, respectively.

**Figure 2.**
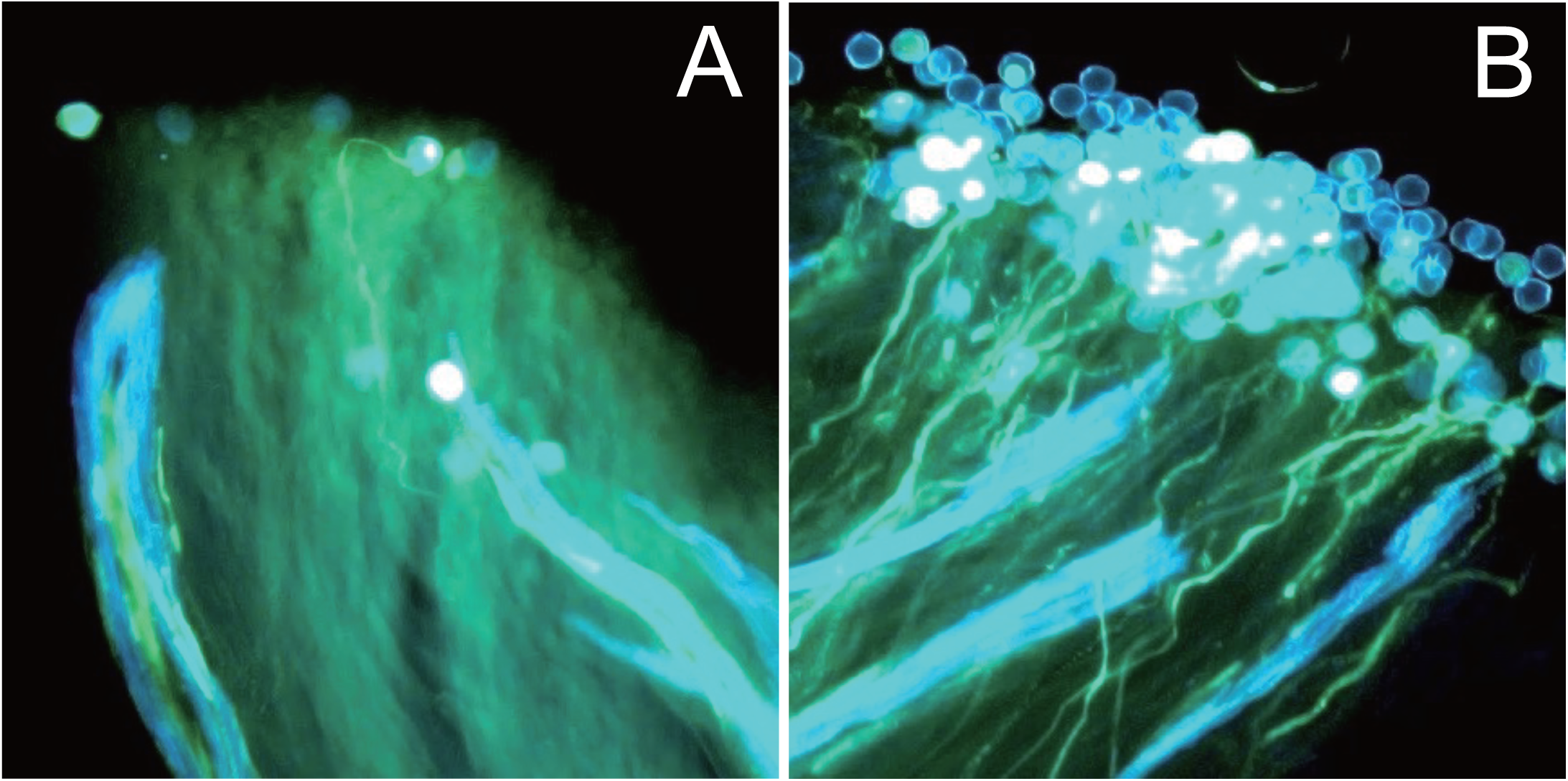
Pollen tube growth on tomato stigmas. Pollen grains and pollen tubes on the stigmas of the transgenic tomato lines (A) *SPI01g07345* and (B) *SPI01g07485*. Images show pollen stained with aniline blue 24 h after pollination.

### *RF2 in* S. pimpinellifolium *LA1670*

Of the F_2_ progeny derived from the cross CMS[MSA1] × LA1670 (Supplementary Figure S1A), two plants (F2_#52 and F2_#53) showed male fertility; both these F_2_ plants lacked the functional *RF1* allele from LA1670. To confirm the inheritance pattern of fertility restoration, the two fertile F_2_ plants were crossed with Dwarf CMS[P] (Supplementary Figure S1A). These crosses produced two fertile F_1_ plants (#52_08 and #53_08), neither of which contained the LA1670 allele. This finding indicated the presence of another novel *RF* locus that was distinct from *RF1*.

To identify this novel *RF* locus, henceforth called *RF2*, plant #52_08 was crossed with Dwarf CMS[P]; this cross produced 13 F_1_ plants (Supplementary Figure S1A). In addition, plant F2_#52 was crossed with MT; this cross produced 36 F_1_ plants (Supplementary Figure S1A). As expected, all progeny from the cross F2_#52 × Dwarf CMS[P] were fertile, but the progeny from the cross F2_#52 × MT segregated into 28 fertile to 8 sterile plants. Of the 49 F_1_ plants produced by these two crosses, 13 fertile and 8 sterile plants were selected for bulked segregant analysis. Sequence reads obtained from the genomic DNA of the bulked sterile and bulked fertile plants were mapped onto the genomic sequence of *S. pimpinellifolium*, LA1670. This identified a 10-Mb *RF2* candidate region that lay between approximately 40.0 and 49.8 Mb of chromosome 2 (Figure 1, Supplementary Figure S2).

### *RF1a2 in* S. lycopersicum *var.* cerasiforme *LA1673*

We tested the mode of fertility restoration in *S. lycopersicum* var. *cerasiforme*, LA1673, using an F_2_ population (n = 153) derived from the cross CMS[P] × LA1673 (Supplementary Figure S1B). All but two F_2_ plants set fruit by self-pollination. Pollen germination was observed, however, in the two plants that did not set fruit, which suggested that *S. lycopersicum* var. *cerasiforme*, LA1673, also possessed the gametophytic mode of pollen fertility restoration.

A BC1F1 population (n = 205) was generated by a cross between an F_1_ hybrid (derived from the cross CMS[MSA1] × LA1673) and O (Supplementary Figure S1C). When the BC1F1 plants had grown to maturity, we observed seed formation and/or pollen germination in 109 plants, but neither fruit setting nor pollen germination were exhibited by the remaining 96 plants, indicating male sterility. The segregation ratio of fertile to sterile plants was close to 1:1 (χ^2^ = 0.824, p = 0.364), indicating that a single gene controlled fertility restoration in this population.

To identify the locus underlying fertility restoration, 106 fertile and 77 sterile BC1F1 plants were subjected to ddRAD-Seq analysis. The obtained reads from fertile and sterile plants were separately merged to generate bulked data and mapped onto the LA1673 genome sequence. This analysis identified an RF locus located in an 80-Mb region of chromosome 1 (Supplementary Figure S2). To identify the RF locus more precisely, fertile BC1F1 plants (female parent) were back-crossed twice with O (male parent) to generate a BC3F1 population (Supplementary Figure S1C). A single fertile BC3F1 plant was self-pollinated to generate a BC3F2 population (n = 31). In addition, two fertile plants, #7 and #20, were emasculated and pollinated using CMS[MSA1] as a pollen donor, producing nine and 19 BC4F1 plants, respectively. The genotypes of all 59 plants were analyzed using 10 CAPS markers. Subsequently, 13 recombinant plants were analyzed further using two additional CAPS markers. This analysis indicated that the *RF* locus was located in a 1.1-Mb region between 80.0 and 81.1 Mb of chromosome 1 (Figure 1). This *RF* locus was named *RF1a2*.

### *RF1d in* S. cheesmaniae *LA0166*

We considered that obtaining the genomic sequence of LA0166 would help us identify *RF* genes in LA0166. We therefore assembled 17.5 Gb HiFi reads obtained from LA0166 into 443 contig sequences that had a total length of 835.6 Mb (Supplementary Table S3). The contigs were aligned with the tomato genome, SL4.0, to establish 12 pseudomolecules, i.e., SCH_r1.0.pmol (Supplementary Table S4). A total of 31,356 genes were predicted from the genomic sequence (Supplementary Table S4). Repetitive sequences occupied 73.4% of the genome (Supplementary Table S5). The accuracy of the genome assembly and gene predictions were supported by BUSCO scores of 98.5% and 93.8%, respectively (Supplementary Table S6). A comparative analysis of genomic structures revealed that, although the sequences of the 12 pseudo-chromosomes essentially corresponded with the chromosomes of LA1670 and LA1673 (Figure 1), several chromosome rearrangements were present, as we detected inversions in chromosomes 1, 5, 8, and 12 and translocations in chromosomes 1, 3, 5, 8, 9, 10, and 11.

The *RF1* locus of LA1670 corresponded with a 2.4-Mb region of LA0166 situated between 81.2 and 83.5 Mb of chromosome 1 (Figure 1); a segmental inversion was detected in the region between 83.4 and 84.4 Mb of LA1670. The *RF1* locus of LA1673 also corresponded with a 1.1-Mb region situated between 81.9 Mb and 82.9 Mb of LA0166 (Figure 1). The 2.4-Mb region of LA0166 was tentatively named the *RF1d* locus. It contained five *PPR* gene copies (*SCH1.0ch01.g38930*, *SCH1.0ch01.g39050*, *SCH1.0ch01.g39780*, *SCH1.0ch01.g39990*, and *SCH1.0ch01.g40160*), whereas the *RF1* candidate regions of LA1670 and LA1673 contained three (*SPI01g07030*, *SPI01g07047*, and *SPI01g07068*) and one (*SLYcer01g06793*) *PPR* gene copies, respectively (Figure 1). In accordance with the gene collinearity detected in these regions, three of the *PPR* gene copies (*SCH1.0ch01.g39780*, *SCH1.0ch01.g39990*, *SCH1.0ch01.g40160*) detected in LA0166 may be orthologs of the three copies detected in LA1670. By contrast, two of the three copies were absent from LA1673, although the remaining copy (*SLYcer01g06793*) may be an ortholog of *SCH1.0ch01.g39780*. In addition, LA0166 possessed two additional *PPR* gene copies (*SCH1.0ch01.g38930* and *SCH1.0ch01.g39050*), which were absent from the LA1760 genome.

## Discussion

We identified an *RF* locus in the same region of chromosome 1 of LA1670 and LA1673, two relatives of tomato (Figure 1). As synteny was observed in this region, this *RF* gene may be allelic or orthologous between the two species. We named this region the *RF1* locus. We also identified two functional *RF* genes, *RF1b* and *RF1c*, both of which encoded *PPR*s, in the sequence surrounding the *RF1* locus in LA1670 (Figure 2; Supplementary Table S2), and therefore hypothesized that these *RF* genes might also be located in the corresponding region of chromosome 1 of LA0166 (Figure 1). Another *RF* locus was identified in chromosome 2 of LA1670 and named the *RF2* locus (Figure 1). These results suggested that fertility restoration by tomato relatives resulted from the accumulation of multiple *RF* genes. These genes may be derived from natural variation or spontaneous mutation. Our results suggested that pyramiding the different *RF* genes into a single plant would generate highly efficient *RF* lines. The SNPs linked to the *RF* loci identified in this study (Supplementary Table S1) will be useful DNA markers for marker-assisted selection in breeding programs to produce *RF* lines through gene pyramiding.

*PPRs* have been identified as *RF* genes in many plant species (Toriyama, 2021). In some species, PPRs are tandemly arrayed in the *RF* regions (Geddy and Brown, 2007). Our analysis of transgenic plants showed that two *PPR* genes, *SPI01g07345* and *SPI01g07485*, showed *RF* function; however, as other *PPR* genes were also present in the candidate regions (Figure 1), further studies are required to determine which of these genes are responsible for *RF* function.

Non-*PPR* genes can also function as *RF* genes in plants (Toriyama, 2021). In maize (*Zea mays*), a gene encoding aldehyde dehydrogenase acts as an *RF* gene that restores CMS caused by Texas (T) cytoplasm (Liu *et al*., 2001), although the molecular mechanism remains unclear. In rice (*Oryza sativa*), a gene encoding a glycine-rich protein that degrades *atp6*-*prf79* mRNA, thus reducing PRF79 protein levels, also acts as an *RF* gene to cause Lead Rice-type CMS (Itabashi *et al*., 2011). Another exception in rice is an *RF* gene that encodes RETROGRADE-REGULATED MALE STERILITY (Fujii and Toriyama, 2009), whose expression is upregulated by retrograde signals from *orf307*, a mitochondrial gene that causes Chinese wild rice-type CMS. The possibility that non-*PPR RF* genes are present in tomato should therefore be kept in mind when seeking to identify *RF* genes.

In general, *RF* genes restore male fertility by suppressing products and/or signals from CMS genes (Toriyama, 2021). Further investigations are therefore required to clarify the molecular mechanisms underlying CMS and its restoration. These investigations should consider multiple aspects of gene regulation, including gene copy number, mRNA processing and degradation, inhibition of translation, structural variation in proteins and protein complexes, and detoxification of CMS gene products. A multi-omics approach, including transcriptomic, proteomic, and metabolomic analyses in addition to genomics and genetics, could be used to identify *RF* genes in tomato. The current study is an important advance in the identification of tomato *RF* genes. Its results will be useful in F_1_ seed production using CMS tomato lines, which currently require a high investment of labor and cost due to their need for emasculation and pollination by hand.

## Supplementary data

**Supplementary Table S1.** CAPS markers linked to the *RF1a2* locus.

**Supplementary Table S2.** *PPR* genes located close to the *RF1a1* locus.

**Supplementary Table S3.** Assembly statistics of the LA0166 genome.

**Supplementary Table S4.** Completeness of the genome assembly and gene prediction.

**Supplementary Table S5.** Chromosome-level assembly of the LA0166 genome.

**Supplementary Table S6.** Repetitive sequences in the LA0166 genome.

**Supplementary Figure S1.** Pedigrees of plant materials.

Line names on the left and right sides of crosses indicate the female and male parent, respectively. Arrows and arrows with dashed lines indicate self-pollinated or crossed progeny and individual selection, respectively.

**Supplementary Figure S2.** Bulked segregant analysis by sequencing.

Arrows indicate the chromosomal positions of *RF1a1* (A), *RF2* (B), and *RF1a2* (C).

## Acknowledgements

We thank Y. Kishida, C. Minami, K. Ozawa, H. Tsuruoka, and A. Watanabe (Kazusa DNA Research Institute) and Dr. Y. Osawa and all the technical members of the T-PIRC center at the University of Tsukuba for technical assistance. Computations were partially performed on the NIG supercomputer at ROIS National Institute of Genetics.

## Author contributions

KS and TA conceived the project. YI, IH, KE, SM, KK, HT, AT, KS, and TA collected the data. YI, IH, KE, SM, KK, HT, KS, and TA analyzed and interpreted the data. KS wrote the manuscript with contributions from YI, IH, KE, SM, KK, and HT. All authors read and approved the final manuscript.

## Data availability

Raw read data was deposited in the Sequence Read Archive (SRA) database of the DNA Data Bank of Japan (DDBJ) under the BioProject accession number PRJDB19708. The assembled LA0166 sequences are available at DDBJ (accession numbers AP028935–AP028946).

